# Single-Cell Multimodal Profiling Highlights Persistent Aortic Smooth Muscle Cell Changes in Diabetic Mice Despite Glycemic Control

**DOI:** 10.1101/2025.04.14.648851

**Authors:** Vinay Singh Tanwar, Vajir Malek, Jingyi Wang, Yingjun Luo, Naseeb Kaur Malhi, Hongpan Zhang, Maryam Abdollahi, Linda Lanting, Parijat Senapati, Sadhan Das, Marpadga A Reddy, Chongzhi Zang, Clint L. Miller, Zhen Bouman Chen, Rama Natarajan

## Abstract

**Background:** Type 2 diabetes (T2D) is associated with accelerated vascular complications like hypertension and atherosclerosis. “Phenotypic switching” of vascular smooth muscle cells (SMC), a major driver of these complications, is enhanced in diabetes. Despite adequate glycemic control, SMC dysfunction can persist due to “metabolic memory” of prior hyperglycemia. However, the mechanisms of hyperglycemic memory associated with persistent SMC dysfunction are unclear. Here, leveraging single-cell (sc) multi-omics, we examined the effect of glucose normalization on transcriptomic and epigenomic changes associated with SMC phenotypic transition in T2D mice.

**Methods:** We treated T2D db/db mice with the antidiabetic drug dapagliflozin (DAPA) (db/dbDAPA) or vehicle (db/db), and non-diabetic control db/+ mice with vehicle for 6 weeks. Dissected aortas were subjected to scRNA-seq, scATAC-seq, and spatial transcriptomics (Xenium) to determine single-cell changes in gene expression and chromatin accessibility.

**Results:** DAPA treatment conferred effective glycemic control in db/db mice, with significant reductions in blood glucose/hemoglobin A1c. scRNA and scATAC-seq analysis of aortas identified major cell populations, including SMC, fibroblasts, endothelial and immune cells. SMC were further clustered into 9 subtypes, including contractile and fibromyocyte-like cells. Cell composition analysis revealed decreases in contractile SMC and increases in vascular remodeling associated fibromyocyte-like SMC in db/db versus db/+ mice. Interestingly, DAPA did not reverse diabetes-induced decreases in contractile markers but reversed changes in several fibromyocyte markers in db/db mice. Pseudotime trajectory analysis revealed increased activities of fibromyocyte enriched transcription factors (TFs) during contractile to fibromyocyte transition. Furthermore, increased expression of TFs regulating fibromyocyte phenotype (e.g. *Atf4*, *Bach1*, *Hand2*, *Fosl2*) in db/db were partially reversed by DAPA, whereas reduced contractile TF (*Mef2c*) expression was unchanged. Spatial transcriptomics analysis further mapped aortic cell types within intact aortas and confirmed that DAPA reversed alterations in key fibromyocyte but not contractile genes in db/db mouse aortas.

**Conclusions:** Persistent epigenetic changes may contribute to sustained vascular remodeling and dysfunction in T2D. T2D reduced contractile SMC gene expression and related chromatin accessibility and promoted phenotypic transition to fibromyocytes. These changes are only partially reversed by a widely used antidiabetic drug like DAPA, underscoring the need for more effective therapies that target hyperglycemic memory.

## INTRODUCTION

Diabetes, a chronic metabolic disorder characterized by hyperglycemia, is a global public health burden.^1^ With increasing prevalence, diabetes is associated with accelerated rates of vascular complications including cardiovascular diseases (CVDs), which are among the leading causes of disability and death.^2–4^ Vascular smooth muscle cells (SMC) are the major cellular components of the healthy arterial wall and help maintain vascular tone in response to metabolic and hemodynamic changes. However, several CVD and type 2 diabetes (T2D) risk factors such as hyperglycemia and dyslipidemia can promote phenotypic switching of SMC from quiescent/contractile to synthetic/proliferative states during diabetic CVD initiation and/or progression.^5–7^ Experimental and clinical studies have shown that prior history of hyperglycemia can lead to sustained progression of diabetic vascular complications despite adequate glycemic control, which is partly due to a metabolic or hyperglycemic memory (MetM) phenomenon, or legacy effect.^2,3,7^ Epigenetic mechanisms have also been implicated in the memory of sustained diabetic complications despite glycemic control.^3,5–8^ However, there are still no systematic transcriptomic and epigenomic studies investigating diabetes-induced SMC phenotypic switching and MetM of SMC dysfunction and crosstalk with other cell types. In the present study, leveraging the power of state-of-the-art single-cell multi-omics and spatial transcriptomics, we investigated these phenomena at single cell resolution.

Single-cell and spatial omics analyses, e.g. single-cell RNA sequencing (scRNA-seq), single cell-assay for transposase-accessible chromatin with sequencing (scATAC-seq) and imaging-based spatial transcriptomics, have yielded unbiased and high-resolution characterization of tissue cellular architecture and cell fate dynamics in both healthy and disease states.^9–13^ Recent multi-omics studies have revealed aortic cell-specific transitions and unique pathophysiological hallmarks of vascular diseases, e.g. atherosclerosis and aortic aneurysms.^6,10–15^ Mouse lineage-tracing studies coupled with scRNA-seq revealed that SMC de-differentiate from contractile to synthetic phenotypes, e.g. fibromyocytes, fibrochondrocytes, and macrophage-like (inflammatory) cell types, during atherosclerosis or aortic aneurysm and dissection.^6,14^ Moreover, scATAC-seq profiling suggested that SMC phenotypic switching can be regulated by alterations in chromatin accessibility, i.e., reduced at contractile gene(s) and increased at gene(s) involved in proliferation, extracellular matrix production, and inflammation.^14^ However, despite diabetes being a major accelerator of vascular complications, such vascular cell phenotypic alterations, and mechanistic drivers of these alterations in diabetes remain unexplored at single cell resolution.

To address these knowledge gaps, we performed integrative multimodal single-cell profiling of aortas from diabetic db/db mice to evaluate the transcriptomic and epigenomic changes occurring in diabetes-induced phenotypic switching, and the effect of the antidiabetic drug dapagliflozin [DAPA, a sodium-glucose cotransporter-2 inhibitor (SGLT2i)] on these changes. We identified discrete SMC populations in diabetic db/db versus db/+ control aortas, accompanied by a decline in contractile SMC gene expression and increase in fibromyocyte-like marker gene expression. Such diabetes associated SMC phenotypic switching was associated with increased expression of key transcription factors (TFs) known to drive SMC transition to fibromyocyte-like phenotype, such as ATF4, BACH1, HAND2 and FOSL2.^15^ Notably, glycemic control with DAPA partially reduced the fibromyocyte-like population/markers without reversing the decline in contractile SMC markers. Finally, we adopted the Xenium transcriptome mapping platform to consolidate our findings in the spatial context. Our data revealed that diabetes induces profound SMC phenotypic switching, which may not be efficiently reversed or prevented with one of the most widely used antidiabetic drug. Our findings highlight the need for more effective therapies for the management of diabetes-associated hyperglycemic memory and vascular complications.

## METHODS

### Study Approval

The animal studies were approved by the Institutional Animal Care and Use Committee at City of Hope (IACUC #10002) in accordance with the National Institutes of Health Guide for the Care and Use of Laboratory Animals. The study with human samples was approved by Institutional Review Board of City of Hope (#01046).

Please see the Major Resources Table in the Supplemental Material.

### Animal experiments

Type 2 diabetic (T2D) male db/db mice, i.e. C57BLKS/J mice with a leptin receptor mutation (BKS.Cg-Dock7^m^ +/+ Lepr^db^/J, Strain # 000642) and nondiabetic db/+ heterozygous littermates (10 weeks old) were obtained from the Jackson Laboratory (Bar Harbor, ME). Throughout the experimental period, mice were maintained under standard environmental conditions in a specific pathogen-free facility in the City of Hope vivarium with food and water provided *ad libitum*. Only male db/db mice were studied to avoid redundancy because female mice also develop diabetes and related complications at similar rates as males (phenotype information for BKS-db from the Jackson Laboratory).

After one week of acclimitization/quarantine, db/db mice showing blood glucose levels (BGL) >250 mg/dL were considered as diabetic and included in the study. db/+ littermates served as nondiabetic control. We randomly divided db/db mice into two groups, and treated them daily with either saline (db/db) or DAPA (db/dbDAPA) by oral gavage (n=12 per group) (Figure 1A). Initially, for four weeks we dosed db/db mice with 1 mg/kg/day DAPA, which is equvalent to the clinically recommended starting dose (5 mg/day oral) to reduce hyperglycemia in T2D patients. In the subsequent two weeks of treatment (4-6 weeks), we increased DAPA dose to 3 mg/kg/day for db/db mice to achieve additional glycemic control.^16^ Body weights and BGL were measured weekly. At the end of six weeks of treatment, glucose tolerance tests (GTTs) were performed, mice were euthanized, and blood samples and thoracic aortas collected for further experments.

**Figure 1:**
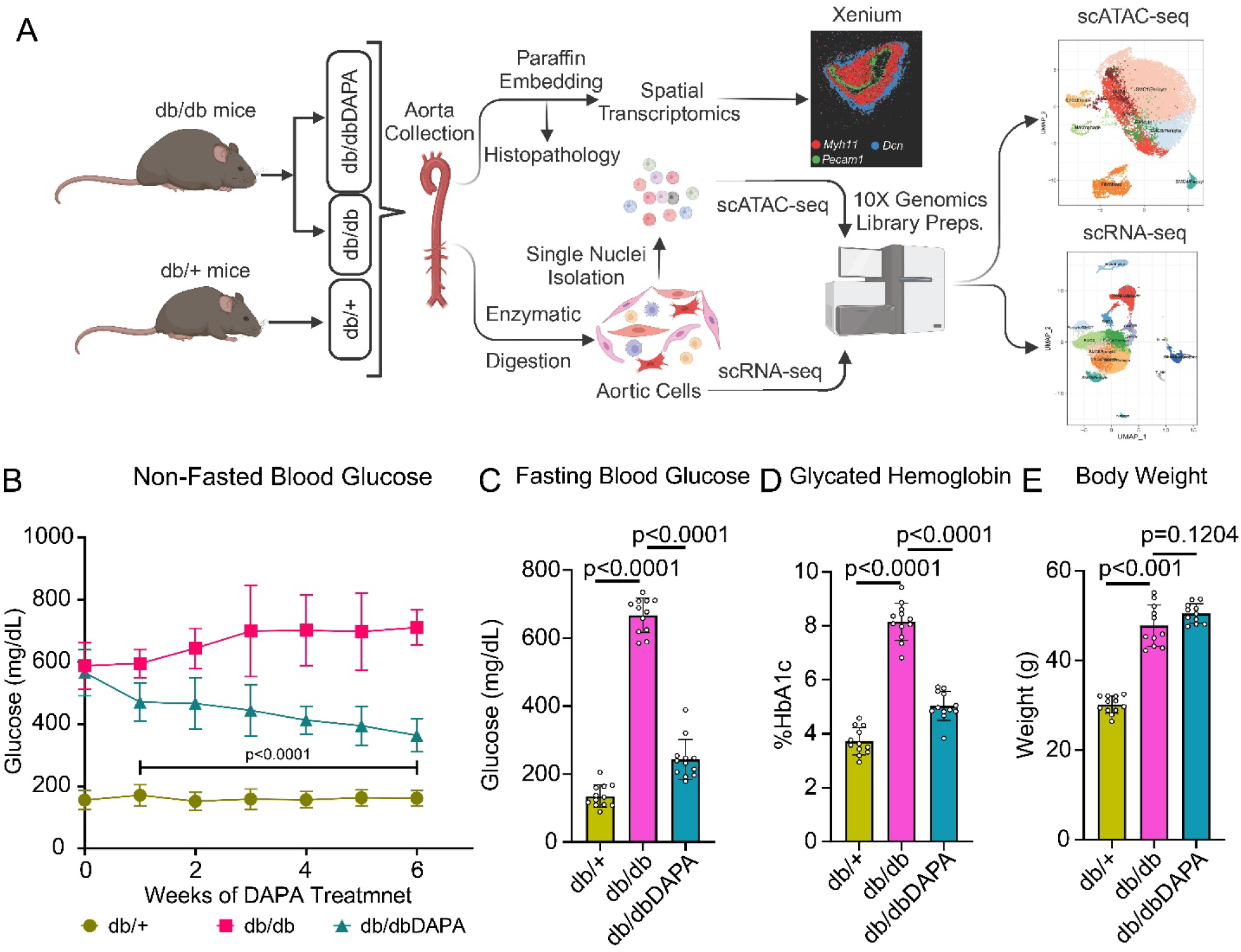
Treatment of diabetic db/db mice with DAPA reduced blood glucose levels and HbA1c. **A.** Schematic of the experimental design. Briefly, type 2 diabetic (T2D) db/db mice were treated with either vehicle (db/db) or dapagliflozin (db/dbDAPA) for six weeks, and their glycemic status monitored. db/+ littermates serve as nondiabetic controls. At the end of six weeks mice were euthanized, and thoracic aortas were collected for further analysis including histopathology, immunohistochemistry (IHC), immunofluorescence (IF), spatial transcriptomics (Xenium), single cell RNA-sequencing (scRNA-Seq), and single cell/nucleus ATAC-sequencing (scATAC-Seq). Details are provided in the Methods section. **B.** Weekly non-fasting blood glucose levels in db/+, db/db and db/dbDAPA mice (n = 12 mice/group). **C-E.** Bar graphs representing fasting blood glucose (**C**), glycated hemoglobin %HbA1c (**D**), and body weight (**E**) in the indicated mice groups at the end of the study (n = 12/group). Data were represented as mean ± standard deviation. Statistical significance and p-values for (**B**) were determined by two-way ANOVA and (**C-E**) by Ordinary one-way ANOVA followed by Tukey’s multiple comparisons tests.

### Measurment of glycemic status and kidney function

Non-fasting (weekly) and fasting (end of the study) BGL were measured from tail vein blood samples using the Alpha Track (Zoetis, MI) glucometer. Hemoglobin A1c (HbA1c) was measured in whole blood samples using mouse HbA1c assay kits (80310, Crystal Chem), and results reported as %HbA1c, as an index of glycemic control. To evaluate kidney function, we assessed surrogate functional markers including albuminuria and albumin-to-creatinine ratio (ACR). We collected 24h urine samples by placing mice in metabolic cages, and measured urine albumin and creatinine levels using an albumin ELISA kit (80630, Crystal Chem) and a pre-validated enzymatic mouse creatinine assay kit (80350, Crystal Chem), respectively.

### Aorta histopathology

Aortas from db/+, db/db, and db/dbDAPA mice were isolated, cleaned and fixed in 10% formalin followed by paraffin embedding and microtome sectioning (5 μm thickness) on glass slides. Deparaffinized tissue slides were stained with hematoxylin and eosin (H&E), periodic acid-Schiff (PAS), or Masson’s trichrome following standardized protocols at the City of Hope Pathology core. All slides were examined by light microscopy in 20x magnification using KEYENCE-BZ-800 series microscope (Keyence, Osaka, Japan). Image analyses were performed using ImageJ software [ImageJ (1.54p)]. H&E staining images were used for the analysis of intima-media thickness. We also assessed extracellular matrix (ECM) accumulation and fibrosis in the tunica media (which contains SMC) by quantifying percentage PAS and trichrome positive area and represented data as fold change over control db/+. For all histopathological image analyses, we used 6 individual measurements for each aorta (proximal region) and 4 aortas per group.

### Single cell preparation from mouse aortas

We pooled aortas from 8 mice into two replicates (4 mice per replicate) for each of the three groups. Aortas (4/group) were cut into ∼1 mm pieces and placed into enzyme mixture containing Liberase DH (1 Unit/aorta, 5401054001, Roche), Elastase (0.125 mg/aorta, E0127, Sigma) and DNase I (Kunitz unit/aorta, 79254, Qiagen) in M199 medium (10060CV, Corning). After incubating at 37 °C for 45 minutes, cell suspensions were filtered through a 70-μm strainer and centrifuged at 500 g for 5 min. The cell pellets were washed with Hank’s balanced salt solution (HBSS) and FACS buffer (HBSS containing 2% FBS) and resuspended in 500 µL of FACS buffer and passed through 35-µm strainer. Cell viability was measured using the Acridine Orange/Propidium Iodide method (Cellometer, Nexcelom Biosciences). Around 10,000 single cells were used for library preparation using the Chromium Next GEM single cell 3’ reagent kits v3.1 (Dual index), Version CG000315 Rev B (10X Genomics) following the manufacturer’s protocol. For scATAC nuclei preparation, cells were washed, counted, and lysed using lysis buffer (10X Genomics) at 4 °C for 5 min. Wash buffer was added after 5 min followed by centrifugation at 500 rcf at 40 °C. Supernatants were removed, and nuclei resuspended in nuclei buffer. Nuclei concentrations were determined using Cellometer. Ten thousand nuclei were processed according to 10X Genomics’ standard protocol using Chromium Next GEM single cell ATAC reagent kits v1.1, Version CG000209 Rev F kit according to the manufacturer’s protocol. Libraries were assessed for quality using TapeStation (Agilent) for fragment sizes ranging from 300-1000 bp, with the peak of fragments around 450 bp. Libraries were multiplexed and sequenced (mean reads per cell (28,000-40,000)) on an Illumina NovaSeq 6000.

### scRNA-seq data analyses

Fastq files from each single cell library were aligned to reference genome mm10 (mm10-2020A) individually using Cell Ranger software (Cellranger v.6.1.1, 10X Genomics). The aligned data sets were analyzed using the R software package Seurat v.3.5.^17^ We filtered out the cells with high mitochondrial reads (<7.5%) and hemoglobin contents (<5%). Potential doublets (individually assessed by counts) were filtered out. Seurat VlnPlot and DimPlot functions were used to generate violin and dot plots. Subset functions were used to create a subset of scRNA-seq data sets. After quality control filtering, we combined all the data sets in one object, followed by normalization, scaling, and principal component analysis. Batch effects were corrected using Harmony (v.1.2.3).^18^ Dimensionality reduction was performed with the RunUMAP command. The FindNeighbors and FindClusters commands were used to cluster the cells from scRNA-seq. UMAP coordinates were extracted from the Seurat object to generate UMAP plots using ggplot2 (v.3.5.1). To gain insights into the cell type composition, mouse aorta scRNA-seq data was integrated with a human atherosclerosis scRNA-seq reference dataset^15^ using FindIntegrationAnchors and IntegrateData commands. Before integration, the gene names from the human atherosclerosis scRNA-seq reference were converted to homologous gene names in the mouse genome using the Mouse Genome Database as a reference.^19^ Cluster labels were transferred from the human atherosclerosis scRNA-seq reference using FindTransferAnchors and then TransferData commands. The cluster labels were refined using positive markers determined using FindMarkers function and other literature-based markers to annotate the scRNA-seq clusters. Gene ontology biological process (GO-BPs) analysis in SMC were performed using differentially expressed genes (DEGs). We selected upregulated genes (adjusted p-values < 0.05, log2FC > 0.25) as positive markers for each subcluster, which were then used to perform GO-BP analysis. Enriched GO-BPs (p-adjust < 0.05) were visualized using dot plots.

### scATAC-seq data analysis

The scATAC-seq data were preprocessed using the 10x Genomics pipeline (Cell Ranger ATAC v.1.2.0) using the mm10 genome and default parameters.^20^ Quality control measurements and filtering were conducted using ArchR (v.1.0.1). Briefly, we retained high-quality cells using the following filters: filterTSS = 4, filterFrags = 1000 and filtered doublets using filterDoublets in ArchR. scATAC-seq reads from all groups were combined at the single-cell level and then mapped to each 500 bp bin across the mm10 reference genome. Quality control metrics (Figure S1A-S1G) indicated similar TSS enrichment (median TSS enrichment = 21.0-22.6) across all groups. Dimensionality reduction was performed using an Iterative LSI based method using the matrix of 500 bp tile-based, with parameters iterations = 2, resolution = 0.2, sampleCells = 10000, n.start = 10 varFeatures = 25000, dimsToUse = 1:30. Clustering was performed using Seurat and clustering resolution =0.8. Clusters were labeled using cell-type annotation from scRNA-seq with addGeneIntegrationMatrix function and normalization parameter was set to “LogNormalize”. UMAP coordinates were extracted from the Seurat object to generate UMAP plots using ggplot2 (v.3.5.1). Marker genes were identified using the getMarkerFeatures function with useMatrix = “GeneScoreMatrix”. The gene score UMAP of known marker genes were generated using plotEmbedding function with quantCut = c (0, 0.99). Cell type specific marker peaks were detected by the getMarkerFeatures function with useMatrix = “PeakMatrix” after peak-calling and plotted using markerHeatmap command. Cell type specific peaks were identified using cut off FDR ≤ 0.05 and Log2FC ≥ 1.5. Motif deviation analysis was performed with chromVAR (v.1.28.0). Variability plots were created with getVarDeviations and density plots with plotGroups. Differential accessible peaks tests between SMC1 and SMC5 were performed using getMarkerFeatures. Volcano plots were generated with plotMarkers (cutOff = FDR <= 0.05 & abs(Log2FC) => 1). Motif enrichments on differential accessible peaks were performed using the peakAnnoEnrichment function.

### Pseudotime analysis

scRNA-seq pseudotime analysis was performed using Monocle3 software in default parameters.^21^ Briefly, we assigned Seurat combined objects into SMC1, SMC2, SMC3, SMC5, SMC8 and SMC10. For scATAC-seq pseudotime analysis, trajectories were created from SMC1, SMC3 and SMC5 using addTrajectory function in ArchR. Heatmaps of features like peaks, motifs, gene expression were generated using plotTrajectoryHeatmap function. Integrated pseudotime analyses heatmaps of peaks, motifs, and genes were generated using correlateTrajectories function in ArchR.

### Xenium data analysis

Xenium spatial transcriptomics experiments were performed on aortas from db/db and db/dbDAPA groups (n = 4/group). We selected 479 genes based on DEG analysis from our scRNA-seq data and literature documenting cell-type markers in the mouse aorta.^6,15,22^ Aortas were cut (5 µM thickness) and transferred to the Xenium slide for further processing at the Integrative Genomics (IGC) core at City of Hope according to manufacturer’s recommendations. Slides were scanned using a Xenium Analyzer. Data sets were analyzed using Seurat (v.5.1.1). Quality control analysis revealed a median of 238 transcripts detected per cell, with more than 10,000 total cells detected across samples. We used Seurat to integrate data from all 8 aortas, followed by UMAP-based clustering to analyze the spatial transcriptomics datasets and could identify all major cell types present in the aorta, including SMC, fibroblasts, endothelial cells, and macrophages.

### Gene ontology enrichment analysis

Gene ontology (GO) biological process enrichment analysis was performed using clusterProfiler with default commands in R. Cell specific GO-BPs were selected from top 20 enriched GO-BPs based on adjusted p-values and plotted as dot plots.

### Human mesenteric scRNA-seq analysis

Cell clusters were generated and annotated using markers based on existing literature and PanglaoDB.^6,22,23^ The scRNA-seq data was analyzed as described.^24^ Differential expression analysis was performed on UMI counts that had been normalized using “LogNormalize” and scaled. The Seurat default, non-parametric Wilcoxon test, was used to analyze genes expressed in at least 10% of cells. The log foldchange (FC) of the average expression was thresholded at 0.25 and a pseudocount of 1 was added to the averaged expression values for differential gene expression analyses.

### Statistical analysis

Data are presented as mean ± standard deviation (SD) (n = number of biological replicates). Prior to comparing between groups, normal distribution of each sample group was confirmed using Shapiro-Wilk test. Statistical significance was calculated using either one-way or two-way ANOVA followed by Tukey’s multiple comparisons tests using GraphPad Prism software (10.0 or above) and p values <0.05 were considered signficant. Bar and XY-plots were created using GraphPad Prism software. Statistical significance is indicated by absolute p-values.

### Data Availability

The authors declare that FASTQ files and matrices from mouse sc-seq [scRNA-Seq, scATAC-Seq, and spatial transcriptomics (Xenium)], human atherosclerosis sc-RNA-seq and sc-ATAC-seq datasets, and human mesenteric artery sc-RNA-seq datasets, have all been deposited to the Gene Expression Omnibus (GEO) database. Accession Codes are available upon request. Mesenteric artery donors were selected based upon the Integrated Islet Distribution Program criteria, with a diagnosis of T2D based on the donors’ medical records as well as the HbA1c of 6.5% or higher. Other supporting data are available within the article and online Supplementary files.

## RESULTS

### DAPA treatment improved glycemic control in db/db mice

We treated T2D db/db mice with DAPA (db/dbDAPA) or vehicle (db/db) for up to six weeks, and non-diabetic db/+ treated with vehicle (db/+) were used as controls (Figure 1A). Diabetic db/db mice displayed significantly higher BGL and body weights (obesity) versus non-diabetic (db/+) mice throughout the study. These elevated BGL were reduced in db/dbDAPA group and continued to decline as the treatment progressed (Figure 1B). At the end of the study, the elevated fasting BGL and HbA1c levels, both parameters of hyperglycemia/diabetes, in db/db mice were nearly normalized by DAPA in the db/dbDAPA group (Figure 1C and 1D), without significant impact on body weight (Figure 1E). SGLT2i, including DAPA, are clinically proven to attenuate diabetic kidney disease (DKD).^25^ As expected, db/db mice showed increased albuminuria and ACR, which were significantly reduced by DAPA (Figure S2A and S2B), confirming the attenuation of diabetes-induced kidney dysfunction by DAPA.DAPA treatment partially attenuates diabetes-induced architectural changes in the aorta.

We performed histopathological assessments of aortic sections to examine the impact of DAPA treatment on diabetes-induced morphological changes in aorta. H&E, PAS and Trichrome staining revealed significant increases in intima-media thickness, extracellular matrix (ECM) accumulation (PAS positive area) and fibrosis (Trichrome positive area) respectively in db/db vs db/+ mice aorta (Figure S3A-S3F). DAPA treatment partially, but statistically significantly, reduced ECM accumulation and fibrosis, with no change in aortic wall thickness when compared to vehicle treated db/db mice (Figure S3A-S3F).

### ScRNA-Seq analysis of mouse aortas from diabetic db/db mice and control db/+ mice identified all major cell types

To elucidate the heterogeneity and phenotypic changes in aortic cells, especially SMC, in diabetes and the effect of glucose normalization, single-cell suspensions were prepared from the aortas of db+, db/db, and db/dbDAPA mice and subjected to scRNA-Seq analysis (Figure 1A). We profiled a total of 56,706 whole aortic cells from all groups (Figure 2A, Table S1). After applying quality control filtering of scRNA-Seq data, cell-based analysis was performed using Seurat. We integrated cells from all 3 groups and performed clustering analysis on a combined Seurat object, which identified all the major cell types previously reported in scRNA-Seq studies using mouse aortas (Figure 2A).^6,15,22^ Using a cell specific marker-based approach, we identified major cell clusters, including SMC, fibroblasts, endothelial cells, macrophages, and other immune cells (Figure 2B and 2C, Figure S4).

**Figure 2:**
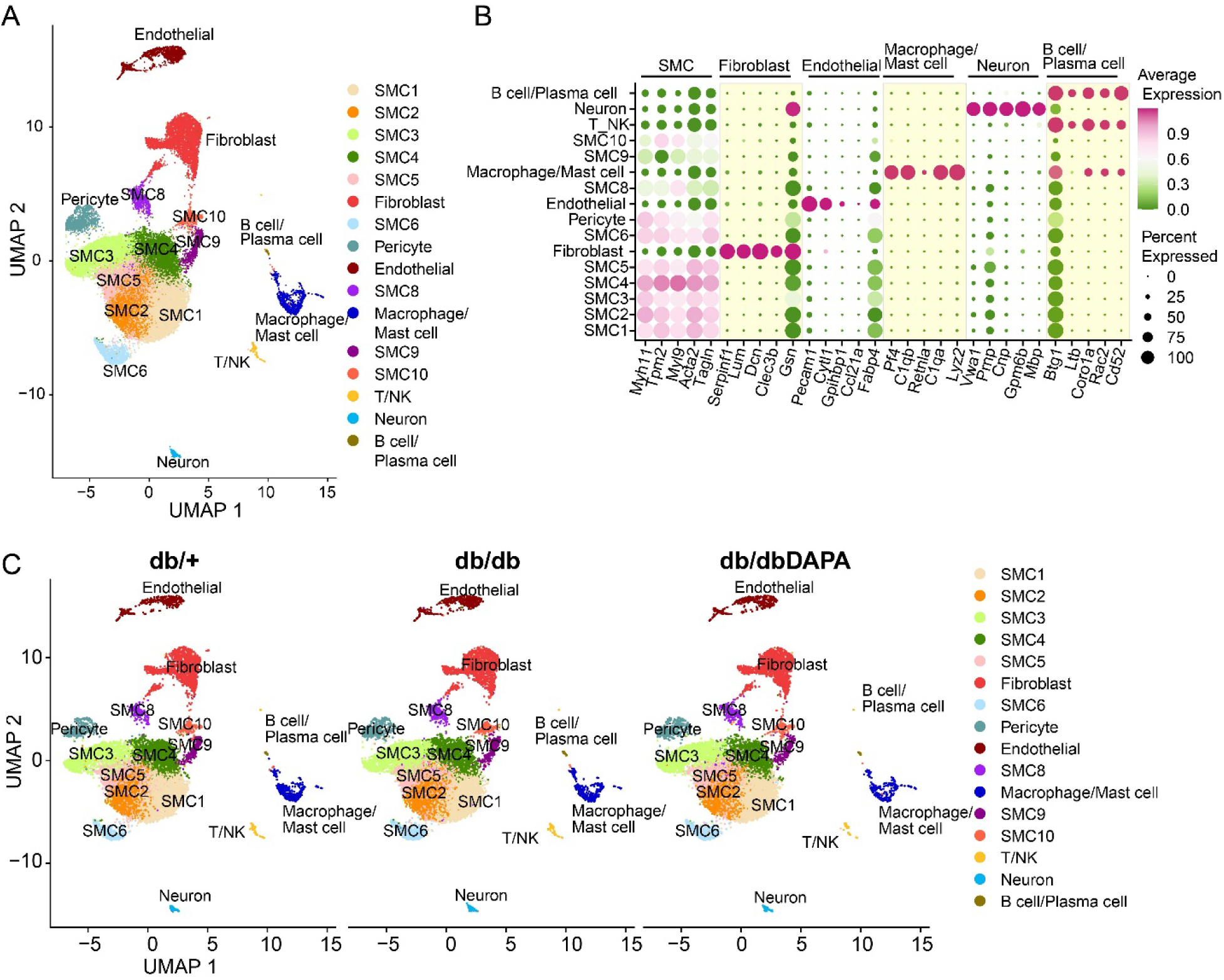
scRNA-Seq analysis reveals aortic cell heterogeneity in control (db/+) and db/db mice treated with vehicle or DAPA treatment. Mice treated as in Figure 1A were subjected to scRNA-Seq. Aortas from 8 mice per group were pooled into two replicates (4 aortas pooled in each replicate) for scRNA-Seq. **A.** Combined UMAP plot of scRNA-Seq showing cellular heterogeneity of aortas from control (db/+), diabetic (db/db) and db/db mice treated with DAPA (db/dbDAPA). Each dot represents a single cell and color representation is based on specific clusters. **B.** Dot plot showing the cell type specific markers for all major cell types identified from scRNA-Seq. The top 5 cell-specific markers are shown. **C.** UMAP of scRNA-Seq data of db/+, db/db and db/dbDAPA mice.

To further characterize the phenotypes/functions of SMC subclusters, we analyzed DEGs after comparing each SMC subcluster with all others. We identified three main SMC phenotypes: contractile SMC (SMC1, SMC2, SMC4, SMC6, SMC9), intermediate phenotype SMC (SMC3, SMC5), and fibromyocyte-like cells (SMC8, SMC10) (Figure 3A-E, Figure S5A-D). Contractile SMC were enriched for BPs such as muscle contraction, muscle cell differentiation, muscle cell development and actomyosin structure organization (Figure 3A and 3B, Figure S5A-C). Intermediate SMC showed enrichment for ECM organization, cartilage development, and collagen fibril organization (Figure 3C and 3D), processes also observed in fibromyocyte-like cells. Fibromyocyte-like cells (SMC8 and SMC10) were enriched in ECM organization, transforming growth factor signaling, bone development, cartilage development, and chondrocyte differentiation, processes known to increase in disease phenotypes as previously reported (Figure 3E, Figure S5D).^12,26,27^ These results confirm the known heterogeneity of aortic SMC subtypes which play important roles in vascular remodeling.

**Figure 3:**
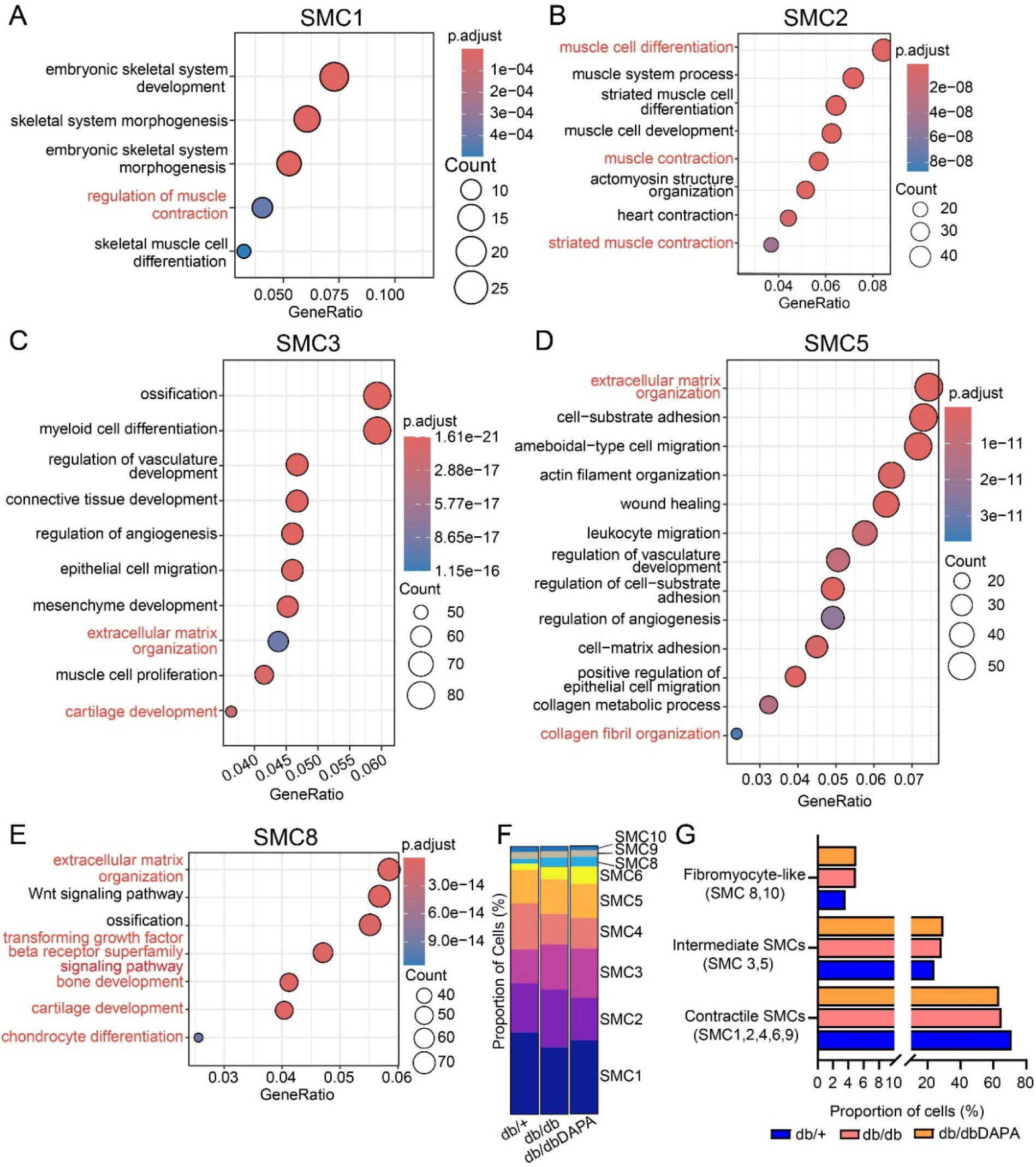
Gene ontology biological processes (GO-BPs) of SMC subtypes in diabetes. **A-E**. Dot plots showing the top enriched GO-BPs in SMC. Each SMC cluster was compared to all other SMC clusters to obtain the upregulated (positive) genes followed by GO-BPs analysis. Top enriched GO-BPs were shown for the indicated SMC. **F.** Stacked bar representation of cell proportions (%) of each SMC cluster in scRNA Seq data in each of the three mice groups. **G.** Bar graph showing the proportion of SMC subtypes (contractile, intermediate and fibromyocyte-like) in db/+, db/db and db/dbDAPA groups. GO-BPs text in red font represent those highlighted in the manuscript text.

### Diabetes induced changes in some SMC composition/proportions are not reversed by DAPA

Next, we performed cell proportion analysis to identify diabetes-induced changes in SMC subcluster proportions across the groups (Figure 3F). The percentage of contractile SMC (SMC1, 2, 4, 6, and 9) was decreased in diabetic mice, and DAPA treatment did not reverse this change (Figure 3G). Notably, fibromyocyte-like cells (SMC8 and 10) were more abundant in both diabetic mice groups compared to control mice (Figure 3G). Intermediate SMC (SMC3 and 5) were increased in diabetic mice and remained increased despite DAPA treatment (Figure 3G). These results suggest that the proportion of contractile SMC are decreased in diabetes and not reversed by DAPA treatment, whereas fibromyocyte-like cells are increased in diabetes and remain increased after DAPA treatment.

### Contractile SMCs markers are not affected by DAPA treatment, whereas some fibromyocyte markers are reversed

Evidence suggests that SMCs undergo transitions into cell types like fibromyocytes and fibrochondrocytes in atherosclerosis.^12,15^ We analyzed SMC dedifferentiation (integrated data sets) using pseudotime analysis of SMC clusters (SMC1, SMC2, SMC3, SMC4, SMC5, and SMC8) with Monocle3, using SMC1 population as starting point (Figure S6). We found that SMC transitioned from SMC1 and SMC2, branching into SMC4 and SMC5, with SMC5 further branching into SMC3 and SMC8. This suggests that, over pseudotime, contractile SMC transition to intermediate forms (SMC3 and SMC5) or fibromyocyte-like SMC (SMC8).

Parallel changes in gene expression were observed during these transitions, including markers for contractile and fibromyocyte-like states. We analyzed the DEGs in SMC5 (intermediate SMC), comparing the two diabetic groups (db/db and db/dbDAPA) to the db/+ group. The heatmap of SMC5 DEGs in the db/db group revealed changes in gene expression vs db/+, and most of these SMC5 gene changes remained unchanged in the db/dbDAPA group (Figure 4A). Furthermore, examination of the GO-BPs associated with the upregulated genes in the diabetic groups (i.e without and with DAPA) revealed an increase in processes related to muscle cell differentiation, transforming growth factor beta (TGF-β) receptor superfamily signaling, ECM organization, and cartilage development (Figure 4B). To further determine if some of these processes are attenuated by DAPA, we compared the SMC5 DEGs in the db/db versus db/dbDAPA group (Figure 4C and 4D). The heatmap revealed a reduction in processes related to muscle cell migration and differentiation, and the TGF-β receptor superfamily signaling, suggesting a protective effect of DAPA treatment on some disease-related GO-BPs (Figure 4D). Next, we compared the DEGs in SMC8 population in the db/db groups versus db/+ group. In the diabetic groups (without and with DAPA), genes associated with muscle cell differentiation, myeloid cell differentiation, and toll-like receptor 4 (TLR4) signaling were upregulated, suggesting these pathways remain dysregulated even after DAPA treatment (Figure 4E and 4F). By comparing the db/db to the db/dbDAPA group in SMC8, we observed a downregulation of genes such as *Apoe* and *Ccn2* indicating some protective effects of DAPA treatment on this gene set (Figure 4G). Together, these data show that DAPA treatment can reverse many, but not all diabetes induced changes in key genes in the indicated SMC subtypes.

**Figure 4:**
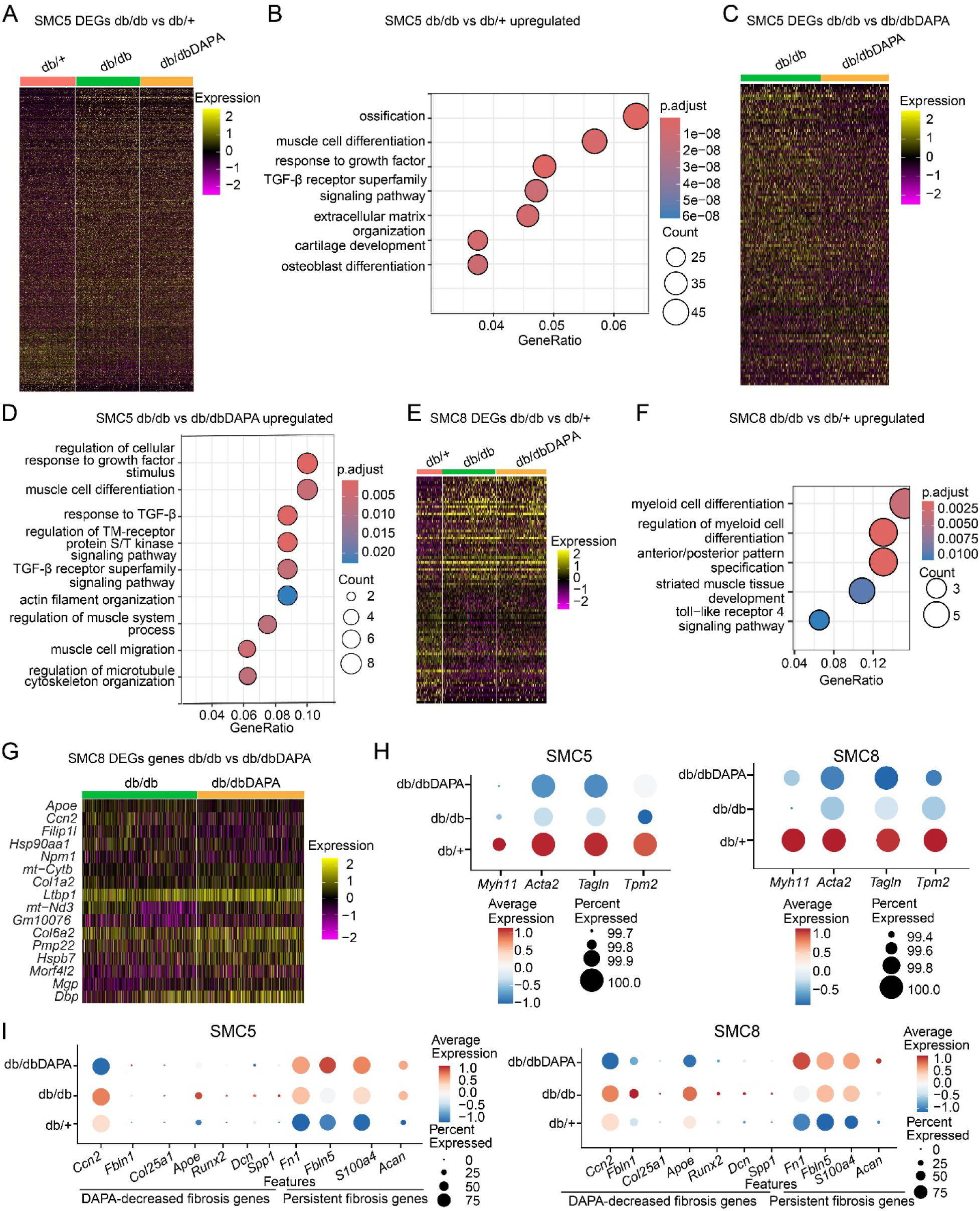
Impact of DAPA treatment on genes and pathways regulated in SMC5 and SMC8 subtypes in db/db mice. **A**. Heatmap of differentially expressed genes (DEGs) in SMC5 in db/db and db/dbDAPA mice versus db/+ mice. **B**. Dot plot showing the gene ontology biological process (GO-BPs) enriched in upregulated genes in the diabetes group compared to control (db/db vs db/+). **C.** Heatmap of DEGs in SMC5 comparing db/db and dbdbDAPA groups. **D**. Dotplot showing the upregulated GO-BPs in SMC5. **E.** Heatmap of DEGs in SMC8 comparing both db/db groups with db/+ group. **F.** Dot plot showing the GO-BPs upregulated in SMC8. **G.** Heatmap of DEGs in SMC8 comparing db/db and db/dbDAPA groups. **H-I**. Expression of candidate contractile and fibrosis related genes in SMC5 and SMC8 showing the persistence of downregulated contractile genes (**H)** and key upregulated fibrosis genes despite DAPA treatment, and other upregulated fibrosis genes that are reduced by DAPA (**I**).

Next, we compared the expression of contractile and fibrosis associated genes in SMC5 and SMC8 and found that SMC contractile genes (*Myh11*, *Acta2*, *Tagln* and *Tpm2*) were decreased in diabetes (db/db vs db/+) and remained decreased even after DAPA treatment (db/dbDAPA vs db/+) (Figure 4H). The expression of key fibrosis and related genes (*Ccn2*, *Fbln1*, *Col25a1*, *Apoe*, *Runx2*, *Dcn* and *Spp1*) were increased in the db/db group and decreased by DAPA treatment. On the other hand, other fibrotic, (*Fn1* and *Fbln5),* inflammatory (*S100a4),* and chondrogenic (*Acan*) genes that were increased in diabetes remained persistently increased after DAPA treatment (Figure 4I). These data show that DAPA treatment can partially reverse increases in some fibrosis markers, whereas, other changes, such as decreased expression of contractile markers and increased expression of key fibrotic inflammatory and chondrogenic genes persisted even after glucose control with DAPA in the db/dbDAPA group. These data support the presence of hyperglycemic/metabolic memory of diabetes on key SMC markers related to vascular dysfunction.

### Fibroblast and endothelial dysfunction markers are increased in diabetes and persist even after DAPA treatment

We next examined the effects of diabetes and glucose normalization on arterial fibroblast, endothelial, and macrophage populations, which also have well-documented contributions to CVD progression.^15^ Using PROGENy (Pathway Resp Onsive GENes),^28^ a tool that infers pathway activity from single-cell transcriptomic data, we found that fibrosis (e.g., TGF-β and WNT) and activities of inflammation-related (e.g., TNF-α and NF-κB) pathways were increased in db/db mice and remained increased following DAPA treatment (Figure S7A). Given that fibrosis in the fibroblast population is a hallmark of disease progression,^29^ we also observed that the increased expression of fibrosis marker genes (*Lum*, *Fbln2* and *Fbln5*) in diabetes continued to be upregulated after DAPA treatment, supporting a memory of diabetes in fibroblasts (Figure S7B).

Because endothelial dysfunction is a hallmark of the initiation of vascular disease, we evaluated endothelial marker and dysfunction-related genes across all three groups. Genes associated with endothelial dysfunction (e.g., *Icam1*, *Vcam1* and *Vwf*) were upregulated in the db/db untreated diabetic group and remained elevated even after DAPA treatment (Figure S7C). Interestingly, marker genes of endothelial homeostasis, such as *Nos3* and *Thbd,* and the endothelial lineage-dependent TF Krüppel-like factor 2 (*Klf2)*,^30,31^ were downregulated in diabetes, which persisted after DAPA treatment. This suggests a persistence of endothelial dysfunction in diabetes despite glucose control, possibly due to metabolic memory.

Analysis of the macrophage population revealed that inflammatory (M1-like macrophage) markers such as *Csf1r*, *Il6* and *Trem1* were increased in diabetes, while anti-inflammatory (M2-like) markers such as *Cd36*, and *Tgfb1* were decreased, which can be associated with the inflammatory macrophage phenotype in diabetes. Interestingly, some inflammatory markers such as *Il6* and *Csf1r* and anti-inflammatory markers *Cd36* and *Tgfb1* were partially reversed by DAPA, suggesting protective effects of DAPA on macrophage inflammatory phenotype (Figure S7D).

### scATAC-Seq identified all major aortic cell types in control and diabetic mice aortas

To identify candidate transcriptional regulators of SMC heterogeneity and phenotypic/epigenetic changes in diabetes, and the effects of glucose normalization, we examined chromatin accessibility using scATAC-Seq across the three groups. Aortas from four mice were pooled per group, and nuclei were prepared from single cells isolated from the db/+, db/db, and db/dbDAPA groups. Using a label transfer approach, we transferred cluster labels from scRNA-Seq to scATAC-Seq, identifying major cell types within the aorta, including five clusters of SMCs, pericytes, fibroblasts, endothelial cells, and macrophages (Figure 5A). Cell-specific markers were used to confirm the identity of these clusters, including *Myh11* for SMC, *Lum* for fibroblasts, *Pecam1* for endothelial cells, and *C1qa* for macrophages, all visualized as peak matrix signals (Figure 5B). Furthermore, UMAP embeddings of the integrated single-cell datasets were generated to visualize the clustering of the combined data (Figure 5A) and explore the effects of diabetes and DAPA treatment (Figure 5C-E). Cell proportion analysis of scATAC-Seq data revealed clear changes in SMC subtypes in diabetes (db/db vs db/+) and with DAPA treatment (db/db vs db/dbDAPA) (Figure 5F). Consistent with the scRNA-Seq data, contractile SMC (SMC1) were reduced in diabetes and remained decreased after DAPA treatment, whereas myofibroblast-like SMC (SMC5) were increased in diabetes and only partially reversed following DAPA treatment. Diabetes induced increases in SMC8 and pericytes, which were further increased in db/dbDAPA group, whereas fibroblasts were not altered in db/db vs db/+ but were increased in db/dbDAPA group (Figure 5F). Thus, cell proportion analysis using sc-RNA/scATAC-Seq revealed decreases in contractile SMC in the aortas of diabetic mice, which persisted even after glycemic control with DAPA. On the other hand, fibromyocyte-like SMC increased in diabetic mice compared to controls and this was partially reversed by DAPA.

**Figure 5:**
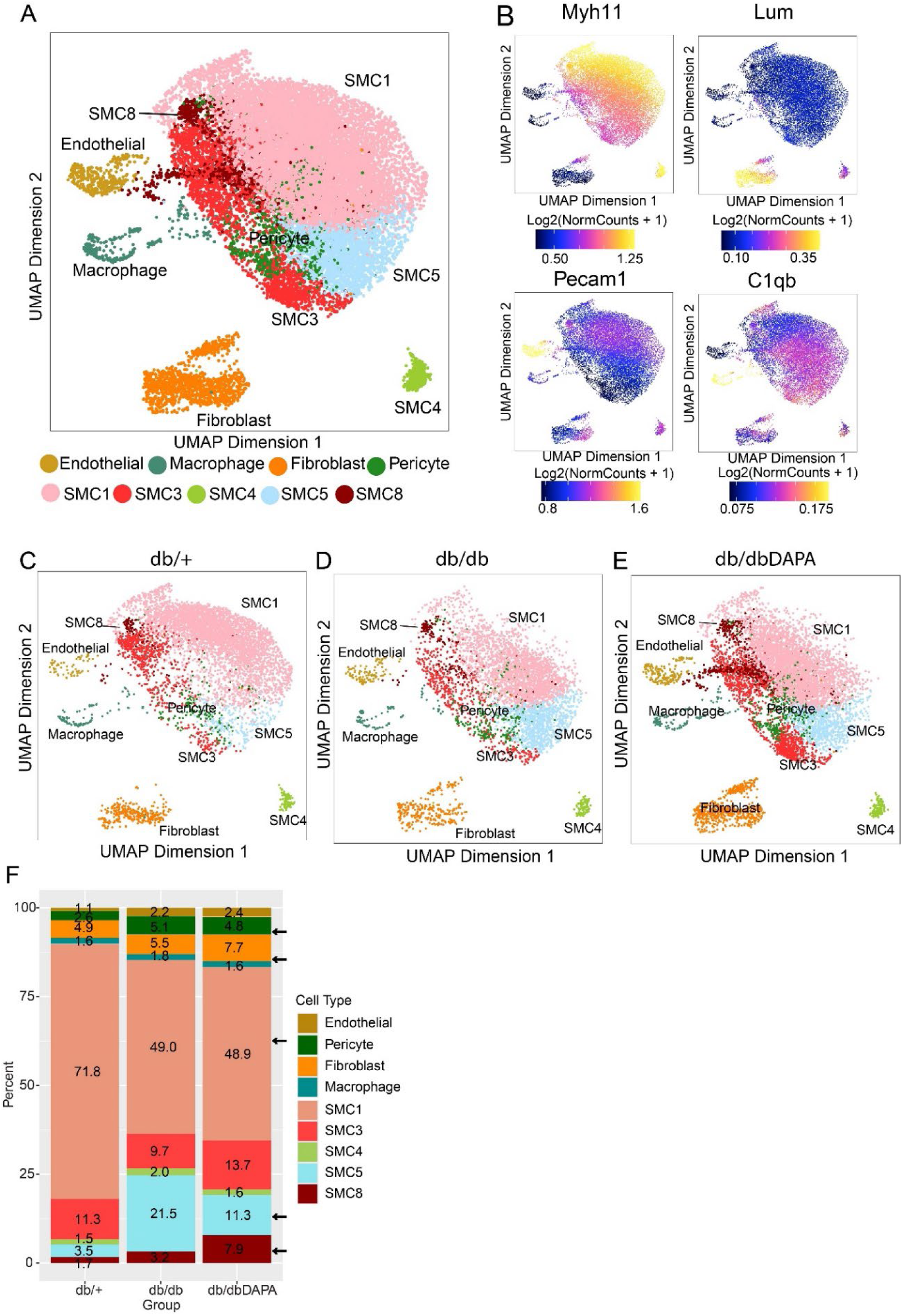
scATAC-Seq identified all major cell types in diabetic aortas. **A.** UMAP and clustering of single-cell chromatin accessibility (scATAC) identified nine clusters of major cell types in diabetes. Each dot represents single cells and clusters based on color. **B.** Specific markers of cell type based on chromatic accessibility. Cell specific markers for SMC (*Myh11*), Fibroblast (*Lum*), endothelial (*Pecam1*) and macrophage (*C1qb*) are shown. **C-E.** UMAP and clustering of scATAC-Seq in control (db/+), diabetic (db/db) and diabetic db/db mice treated with DAPA (db/dbDAPA). **F.** Stacked bar plot showing the percentage of each cell type present in aortas from each group. Colors represent the different cell types in the aorta and the numbers within the bars indicate the percentage of each cell type within each group. Arrows point to those showing major changes in cell proportions in diabetes and DAPA treatment.

Next, we examined cell-type-specific ATAC-Seq peaks (see Methods) and identified 56,423 features across the combined dataset (Figure 6A). TF motif enrichment analysis revealed enrichment of contractile-associated TFs, including MEF2A/B/C/D, SMAD1/5 and TCFAP2D in the SMC1 population (Figure 6B). Notably, fibroblast-associated TFs (BACH1/2, FOS, FOSB, JUNB, JUND, NFE2L2, and SMARCC1) were enriched in SMC3 and SMC5. Among these, fibromyocyte-associated TFs were highly enriched in SMC5, followed by SMC3, supporting their fibromyocyte-like phenotype (Figure 6B). Endothelial TFs, including ERG, ETS1, and ETV2, key regulators of endothelial homeostasis, were enriched in endothelial scATAC-Seq peaks. Macrophage-associated TFs, such as ELF1, SFPI1, SPIC, and ELF5, were enriched in macrophages. Motif deviation analysis in the integrated dataset (Figure 6C) revealed a similar enrichment of these TFs at the single-cell level, as well as increased activity of fibromyocyte-associated TFs in SMC3 and SMC5 (Figure 6C and 6D, Figure S8A-J). Moreover, to identify TF regulators of SMC5 (fibromyocytes) compared to contractile SMC1, we performed differential peak analysis and identified 9,320 upregulated peaks and 692 downregulated peaks in SMC5 vs SMC1 (Figure 6E). Motif enrichment analysis of the upregulated peaks in SMC5 revealed significant enrichment of fibromyocyte-related TFs, including SMARCC1, FOS, BACH1, and BACH2 (Figure 6F), while downregulated peaks showed enrichment of TF motifs (WT1, KLF15 and ZFP219), which are known to regulate SMC contractile gene expression (Figure S8J.

**Figure 6:**
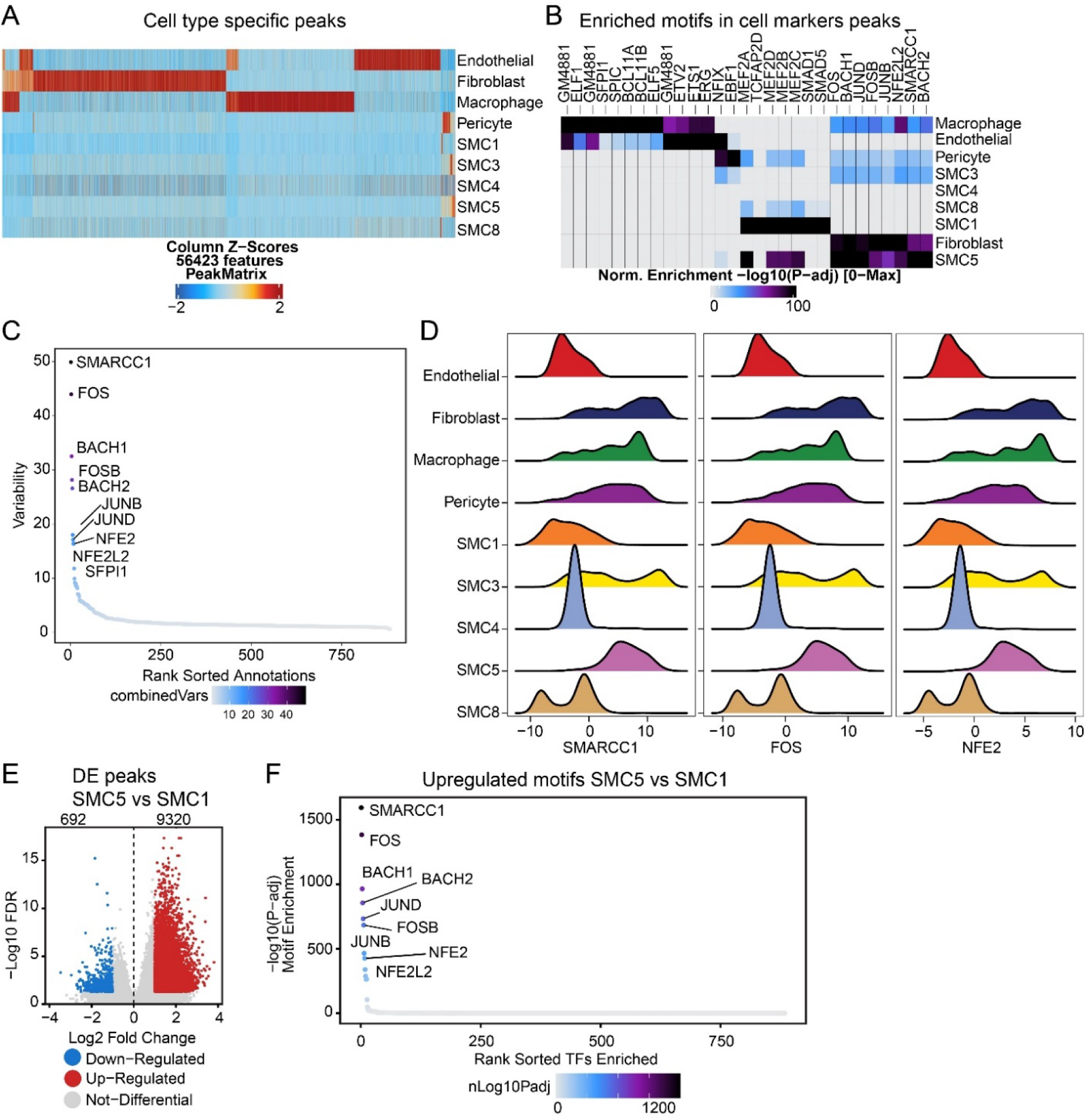
Chromatin accessibility and enrichment of transcription factors (TFs) motifs in all cell types (including fibromyocyte-like cells) in diabetes. **A.** Heatmap of cell specific chromatin accessibility peaks in all the major cell subtypes in diabetes and DAPA treatment. **B.** Heatmap of TF motif enrichment in cell-specific chromatin accessibility peaks. **C.** Ranked variability of TF motifs in chromatin accessibility data in diabetes highlighting the highly variable TFs. **D.** Plot of fibromyocyte-like cell TFs that show high variability in cell-specific chromatin accessibility data sets. SMARCC1 and FOS and NFE2 transcription factors are shown. **E.** Volcano plot of differentially enriched (DE) scATAC-Seq peaks in SMC5 (fibromyocyte-like cells) compared to SMC1 (Contractile SMC). **F.** TFs motifs enriched in upregulated chromatin accessibility peaks in SMC5 (vs SMC1).

Pseudotime trajectory analysis also further verified the data that SMC dedifferentiate into fibromyocyte-like SMC in diabetes and DAPA treatment only partially reverses this transition. To identify the gene regulatory programs and TF motifs associated with the phenotypic transition from contractile SMC to fibromyocytes in diabetes, we performed pseudotime trajectory analysis using ArchR, setting SMC1 (contractile SMC) as the starting point (Figure 7A). The pseudotime trajectory analysis revealed that both the chromatin accessibility score for *Myh11* (contractile marker) and its gene expression (from integration with scRNA-Seq data) decreased during transition from SMC1 to SMC5 (Figure S9A and S9B). In contrast, we observed an increased enrichment of fibromyocyte-related TF motifs such as JUND, JUNB BACH1, BACH2, and FOSB (Figure 7B, bottom heatmap). By integrating scRNA-Seq data, we also observed decreased contractile marker gene expression, such as *Acta2*, *Tagln*, *Myh11*, *Myl6*, and *Tpm2*, and increased expression of fibromyocyte-associated genes such as *Jund*, *Atf3*, *Fosb*, *Fn1*, *Vim*, and *Lgals1* (Figure 7C). Integrated pseudotime analysis using aligned scATAC- and scRNA-Seq data showed contractile SMC TFs, such as MEF2C, with decreased motif enrichment and gene expression, while fibromyocyte-associated TFs, including FOSL2, NFKB1, and SMARCC1, showed increased enrichment and expression during the SMC1 to SMC5 pseudotime transition (Figure 7D and 7E, Figure S9C and S9D).

**Figure 7:**
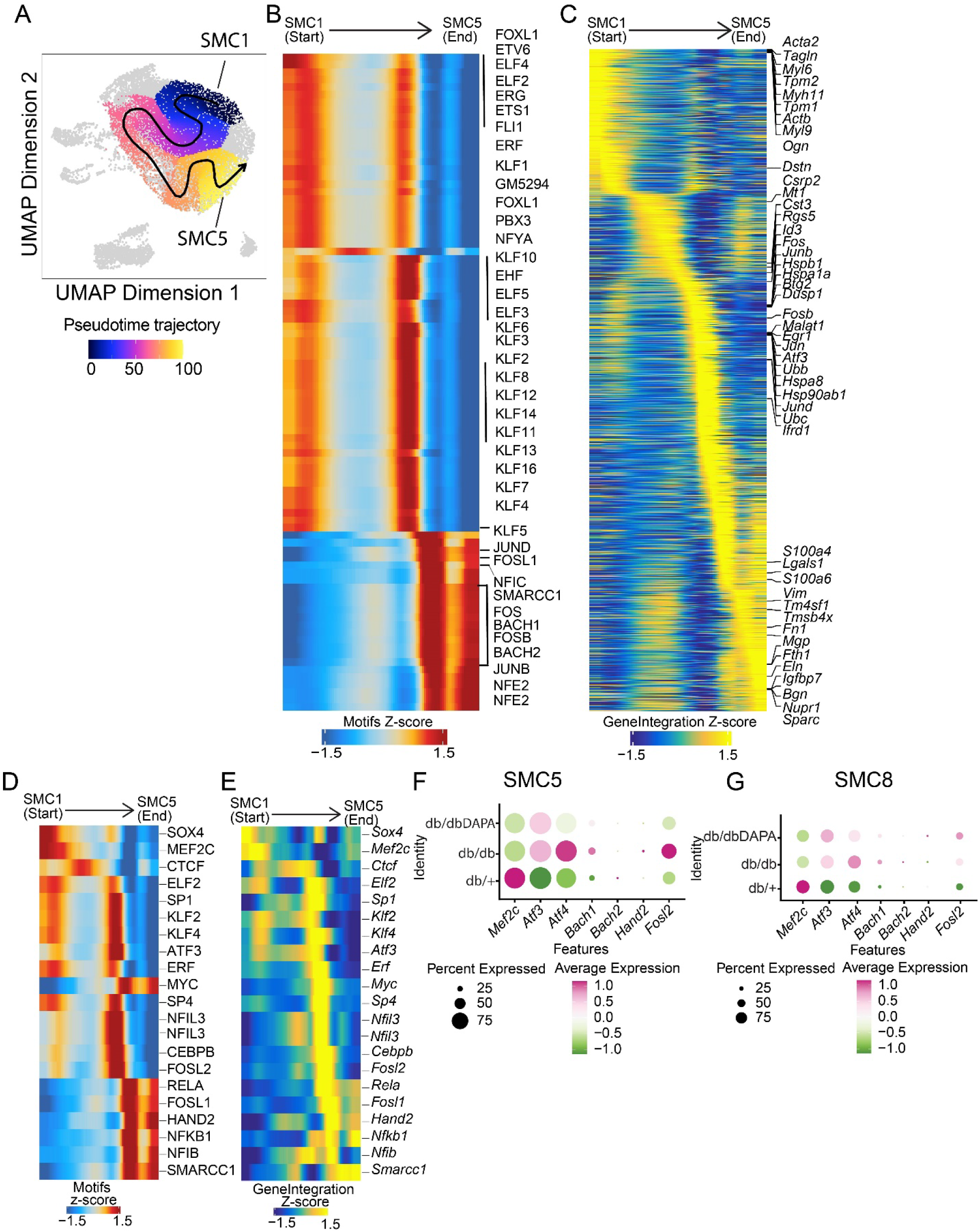
Pseudotime analysis of SMC (SMC1, SMC3 and SMC5) reveals the origin of SMC5 from SMC1. **A.** UMAP showing the pseudotime trajectories based on ATAC-Seq data sets. **B**. Heatmap of indicated TF motif enrichments during pseudotime transition. Values represent motif Z-scores. **C.** Heatmap of integrated gene expression (from scRNA-Seq) during pseudotime transition. **D-E.** Integration of scATAC and scRNA-Seq data reveal pseudotime trajectory during SMC1 to SMC5 transition. Heatmap of pseudotime matched motif from scATAC-Seq and gene expression from scRNA-Seq during SMC1 to SMC5 pseudotime transition. **F-G**. Dot plots showing the gene expression of fibromyocyte related TFs in SMC5 (F) and SMC8 (G).

To further assess the impact of diabetes and DAPA treatment on contractile and fibromyocyte-associated TFs, we analyzed scRNA-Seq data from SMC5 and SMC8 which are fibromyocyte-like cells that showed changes in cell proportion between groups based on scRNA-Seq and scATAC-Seq data sets. We found that contractile TF gene (*Mef2c*) was downregulated in SMC5 in diabetic mice and remained downregulated even after DAPA treatment (Figure 7F). Additionally, expressions of fibromyocyte-associated TFs such as *Atf4*, *Bach1*, *Hand2*, and *Fosl2* were upregulated in diabetic mice, and DAPA treatment only partially inhibited their expression (Figure 7F). Similar findings were observed in SMC8 (Figure 7G), supporting the conclusion that diabetes downregulates contractile TF expression while upregulating fibromyocyte-like TF gene expression in fibromyocyte-like cells (SMC5 and SMC8). This further reinforces the data suggesting that DAPA only partially suppresses disease-related phenotype switching of SMC in diabetes.

### Spatial transcriptomics data confirm sc-Seq data, including non-reversal of contractile genes and partial reversal of fibromyocyte associated genes in DAPA-treated diabetic mice aortas

To assess SMC phenotypic switching in diabetic arteries at single-cell and spatial resolution and to validate our findings from the sc-Seq, we leveraged the state-of-art Xenium targeted *in situ* transcriptomics platform. Using cell-type-specific markers and cluster analysis, we annotated major cell types (e.g., fibroblasts, endothelial cells, and macrophages), as well as SMC subtypes (SMC1 through SMC8/Pericytes) in the mouse aorta (Figure 8A-C). Spatial transcriptomics data demonstrated highly specific expression patterns of cell type-specific genes, marking their positions within the artery sections (Figure 8D and 8E). Contractile marker genes (*Myh11*, *Acta2*, *Tagln* and *Tpm2*) in SMC, which were downregulated in the diabetic db/db group in the scRNA-Seq data, showed even lower expression in the db/dbDAPA group (Figure 8F, 8I and 8J; Figure S10A-E). Fibrosis and chondrogenic markers such as *Fn1* and *Acan* found to be upregulated in db/db group, remained upregulated in the db/dbDAPA group (Figure 8G and 8K; Figure S10F). Some fibromyocyte-associated genes (*Apoe*, *Dcn*, *Spp1*, *Col1a1* and *Col1a2)* that were increased in db/db group were decreased in the db/dbDAPA group (Figure 8H-L, Figure S10F).

**Figure 8:**
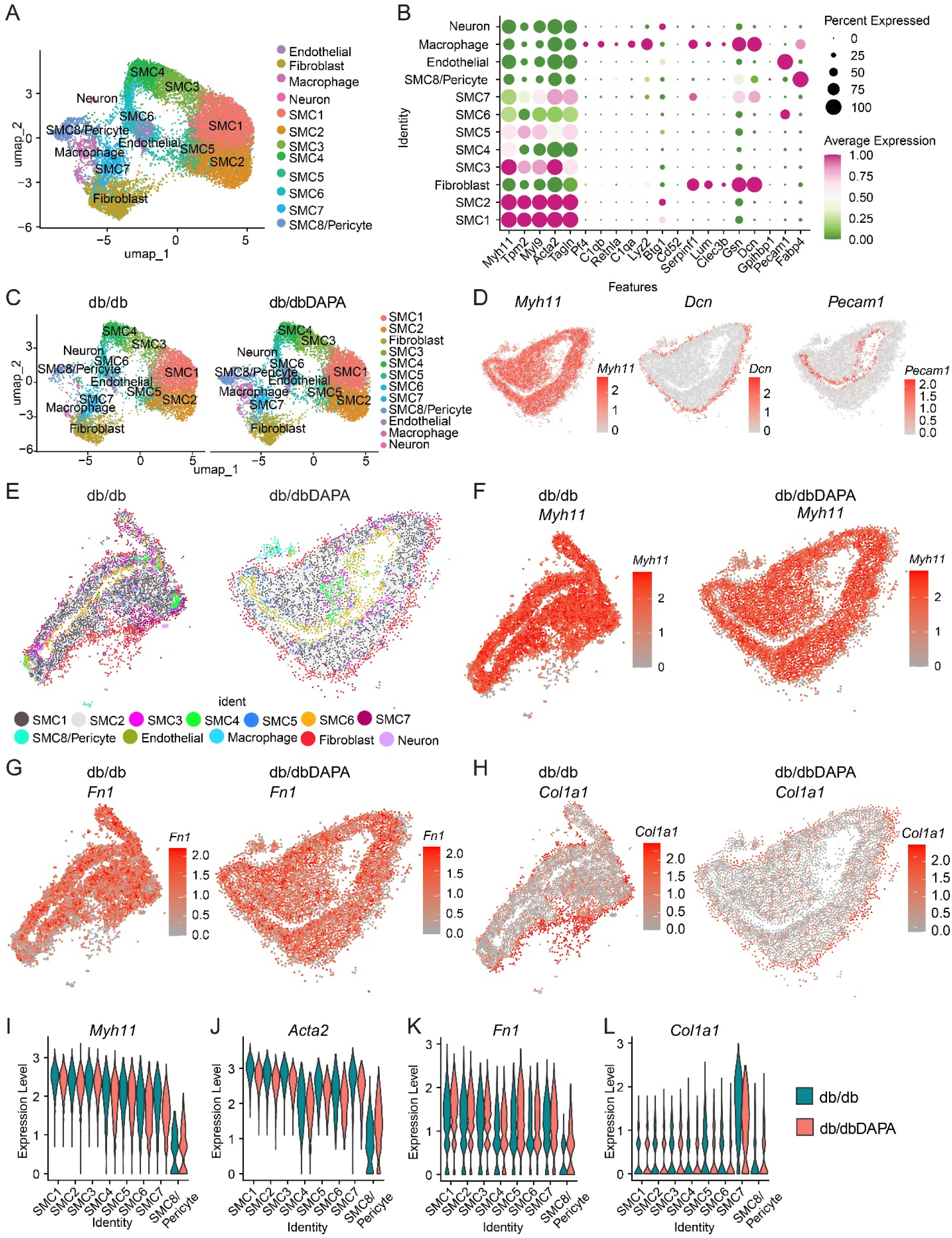
Xenium Spatial Transcriptomics profiling reveals major aortic cell types and organization and validates sc-Seq data. **A.** UMAP clustering of combined Xenium data sets from diabetic db/db and diabetic mice treatment with DAPA (n=4) for 6 weeks. **B.** Dot plot showing the cell specific markers from combined analysis of Xenium data sets. **C.** UMAP plot of clustering of cells type in aorta from db/db and db/dbDAPA mice. **D.** Xenium data showing one of the db/dbDAPA aortas highlighting cell specific markers of SMC (*Myh11*), fibroblast (*Dcn*) and endothelial cells (*Pecam1*). **E.** Aortic cell clusters are depicted clearly in aortic sections in Xenium data sets from db/db and db/dbDAPA group. **F-H**. Representative images of Xenium data from db/db and db/dbDAPA aortas showing the expression of contractile (*Myh11*) and Fibrosis markers (*Fn1 and Col1a1*). **I-L.** Violin plots depicting expression of contractile (*Myh11*, *Acta2*), and fibrosis associated genes (*Fn1, Col1a1*) in db/db and db/dbDAPA groups (showing DAPA reduces *Col1a1* fibrosis marker).

Next, we examined the expression of fibroblast (Figure S11A-D) and EC-specific genes (Figure S11E-G) and found that DAPA treatment decreased the expression of a few fibrosis (*Fbln1* and *Fbln5*) and endothelial dysfunction marker (*Vwf*) suggesting some protective effects in these cells (Figure S11D-G). On the other hand, endothelial function markers *Nos3* and *Klf2* showed even further decreased expression after DAPA treatment (Figure S11G), consistent with the scRNA-Seq data.

### SMC contractile genes are decreased and fibromyocyte genes are increased in human mesenteric artery SMC from donors with pre-diabetes and diabetes

To determine human relevance, we examined scRNA-Seq data obtained from mesenteric arteries obtained from non-diabetic, pre-diabetic, and diabetic donors for the expression of candidate marker genes altered in both db/db and db/dbDAPA groups, especially those genes showing persistent expression in both these mouse groups. Demographic characteristics of the deidentified donors are shown in Table S2 and cell numbers from scRNA-Seq in Table S3. The expressions of SMC contractile genes (*MYH11*, *ACTA2*, *TAGLN* and *TPM2*) were decreased in pre-diabetic and diabetic SMC compared to non-diabetic (non-DM) (Figure S12A), while the fibromyocyte markers (*ACAN*, *S100A4*, *FBLN1*, *FBLN5* and *FN1*) showed increased expression in pre-diabetic or diabetic SMC, further confirming decreased expression of contractile genes, and increased expression of genes related to fibromyocyte transition in human diabetes (Figure S12A).

We also found that the expression of fibrosis associated genes in fibroblasts (*LUM*, *FBLN2*, and *PDGFRA*) was increased (Figure S12B). In ECs, expression of dysfunction markers (e.g., VCAM1, ICAM1, *VWF*, *CXCL1* and *CD36*) was increased whereas endothelial homeostasis markers *NOS3* and *THBD* was decreased in pre-diabetic and diabetic compared to non-DM samples (Figure S12C). Furthermore, inflammatory markers (M1-like macrophage markers, e.g. *CSF1R* and *TREM1*) in macrophages were increased and anti-inflammatory markers (M2-like, e.g. TGFB1 and CD163) were decreased in prediabetes or diabetes compared to non-DM samples (Figure S12D). These results in human samples validate the findings regarding diabetes-related key genes from mouse aortas.

## DISCUSSION

Under diseased states, aortic SMC transdifferentiate into various phenotypes, including fibromyocytes, fibrochondrocytes, and macrophage-like cells ^6,12,15,23^. Fibromyocyte-like cells are characterized by low levels of contractile gene expression (e.g *Myh11*, *Acta2*, *Tpm2* and *Tagln*) and high levels of fibrosis-related genes (e.g. *Fbln1*, *Fbln5*, *Ccn2*), while fibrochondrocytes exhibit low expression of contractile genes and high levels of chondrocyte and cartilage related genes such as aggrecan (*Acan*). Similarly, macrophage-like SMC are characterized by high levels of inflammatory gene expression. Dedifferentiated and transdifferentiated SMC tend to increase in numbers in atherosclerotic disease, contributing to plaque development and instability. However, although SMC dysfunction is known to increase in diabetes, the phenotypic changes in specific SMC subsets and other aortic cells occurring in vivo during diabetes are unclear. Furthermore, the extent to which these cellular transitions persist even after treatment with glucose lowering drugs and whether there is a memory of SMC dysfunctional states are also unclear.

Here, we investigated whether the antidiabetic drug SGLT2i DAPA can fully or partially protect against diabetes induced SMC and other aortic cell heterogeneity/transitions associated with vascular dysfunction. We performed scRNA-Seq, scATAC-Seq and Xenium spatial transcriptomic analysis to identify the aortic cell types, cell-specific genes, TFs and other regulators differentially regulated in aortas of diabetic mice with and without DAPA treatment. Our results show that contractile SMC genes, e.g. *Myh11*, *Acta2*, *Tagln* and *Tpm2,* are decreased in diabetes and DAPA treatment does not reverse this effect especially in aortic SMC5 and SMC8 populations. We further validated our single-cell based results using spatial transcriptomics analysis, which also showed that contractile gene expression remains similar in db/dbDAPA compared to db/db group. In addition, consistent with other single-cell datasets,^15^ we identified a fibromyocyte-like cell population of SMC in the mouse aorta, which was increased in diabetes, and DAPA treatment only partially reversed this effect. DAPA also partially reversed the diabetes-induced expression of TFs that promote fibromyocyte differentiation, further supporting the partial suppression of fibromyocyte-like cells and their phenotype by DAPA treatment in diabetes.

TFs play key roles in SMC phenotypic transitions during vascular remodeling and disease. Prior scATAC-Seq studies in atherosclerotic human coronary arteries have nominated TFs like ATF3, JUN, BACH1, BACH2 enriched in fibromyocyte and fibrochondrocyte-like cells.^12^ Our results highlight similar TFs (JUN, BACH1, BACH2) increased in fibromyocyte-like cells and MEF2C decreased in contractile SMC, supporting a key role for these TFs in SMC trans-differentiation in diabetes. Moreover, we observed increased expression of TFs like *Atf4*, *Bach1*, and *Fosl2* in diabetes which were decreased by DAPA treatment, indicating protective effects of DAPA on these factors. Overall, these results identify transcriptional mechanisms for diabetes-induced SMC phenotypic transitions, which persist during glycemic control.

SGLT2i, originally developed to treat T2D, also demonstrate remarkable cardio- and reno-protective effects in individuals with and without diabetes.^32,33^ The mechanisms underlying beneficial effects of these inhibitors include normalized blood glucose, improved kidney function, and blood pressure control.^32,33^ In this study we confirmed that treatment with the SGLT2i DAPA decreases blood glucose levels, HbA1c, and improves kidney function. Interestingly, our aortic single-cell multi-omic data demonstrate that, while DAPA exerts protective effects in suppressing the fibromyocyte-like phenotype transitions in T2D mice, the persistent loss of SMC contractile phenotype despite DAPA treatment suggests that DAPA may be insufficient to prevent or attenuate persistent vascular dysfunction in diabetic patients. Given that our DAPA treatment was for only 6 weeks, it is possible that a longer treatment duration might have more superior beneficial effects.

EC dysfunction is another hallmark of both atherosclerotic and diabetic vascular diseases. Nitric oxide (NO) plays a critical role in vasodilation, contributing to endothelial homeostasis and protection against endothelial dysfunction. Reduced NO levels are associated with endothelial dysfunction in diabetes and atherosclerosis. Moreover, thrombomodulin (*Thbd)*, which converts thrombin into an anticoagulant, plays a protective role in thrombosis, inflammation, and fibrinolysis.^34–36^ Our single-cell results revealed reduced EC expression levels of *Nos3* and *Thbd,* as well as the TF *Klf2* (which regulates the expression of *Nos3* and *Thbd*) in diabetes, as expected. Interestingly, DAPA treatment failed to restore their expression. In addition, markers of endothelial dysfunction, such as *Vcam1* and *Vwf*, that were elevated in ECs of diabetic mice, were not normalized by DAPA treatment. These data support the presence of T2D induced memory, which may also contribute to persistent EC dysfunction in diabetes.

While our study provides molecular insights into the metabolic memory during diabetes we acknowledge some limitations. We note that larger numbers of profiled cells and aortic tissues in single-cell and spatial studies may be necessary to fully disentangle the impact on SMC-related cell populations, such as pericytes and fibroblasts, as well as macrophages and ECs. It should also be noted that DAPA did not significantly reduce body weight in diabetic mice in the time period tested, and hence some observed irreversible/persistent changes may also be partly attributed to obesity. Future experiments with longer treatment times and other antidiabetic drugs like GLP-1 receptor agonists (which are both antidiabetic and anti-obesogenic) will help determine if changes noted in our study persist despite body weight reduction. Evidence shows that lowering ApoB-containing lipoproteins can reduce SMC-derived fibromyocytes and chondromyocytes from plaques.^37^ Hence it may be worth examining whether ApoB lowering drugs would have additive or synergistic effects with antidiabetic drugs on SMC modulation.

In conclusion, our integrated sc-multi-modal datasets revealed for the first-time diabetes/hyperglycemia induced (epigenetic) memory in the SMC, endothelial cells and fibroblasts with single cell and spatial resolution, which is not entirely reversed by SGLT2i DAPA. The identified cellular and molecular changes are likely responsible for persistent vascular dysfunction in patients with prior uncontrolled diabetes despite subsequent glucose control. Combination therapies including medications that maintain contractile SMC and EC functions might confer more vascular benefits for these patients and improved clinical outcomes with reduced long term persistent vascular complications.

## AUTHOR CONTRIBUTIONS

VST and VM performed animal experiments and wrote the first draft of the paper. JW, VST, HZ, VM, NKM and PS analyzed the data. VST, JW, HZ, VM, YL, NKM, PS, SD contributed reagents/materials/analysis tools. RN, ZBC, MAR, CLM, CZ, JW, YL, NKM edited the manuscript. VM, LL and MA performed animal experiments. VST, VM and LL processed the mouse aortas and prepared single cells for scRNA-Seq, scATAC-Seq and Xenium. YL prepared the sequencing libraries and designed the Xenium gene panel. VM and MA performed histochemistry and microscopy. CLM and CZ provided funding and supervised research secondarily related to the study. RN, and ZBC provided the primary funding and supervised the study. RN conceptualized the work, helped in the experimental design, edited the manuscript, acquired funding, coordinated and supervised the study.

## ACKNOWLEDGEMENTS

We are grateful to Dr. Jinhui Wang and Dr. Xiwei Wu (Integrative Genomics Core, City of Hope) for assistance with the sequencing and spatial transcriptomics profiling, and to Pathology research services at City of Hope. We are grateful to Dr. Suchismita Dey (Department of Diabetes Complications and Metabolism, Beckman Research Institute of City of Hope) for help in cleaning mouse aortas and single cell preparations. Research reported in this publication includes work performed in the Integrative Genomics Core and DNA/RNA Synthesis Core (supported by the National Cancer Institute of the NIH under grant no. P30CA033572), and the Light Microscopy Core of City of Hope.

## SUPPLEMENTAL MATERIAL

- Supplemental Figures S1-S12
- Supplemental Tables S1-S3
- Major Resources Tables

## Novelty and Significance

### What is known?

1. Diabetes is associated with vascular smooth muscle cell (SMC) dysfunction and significantly accelerated vascular complications like atherosclerosis and hypertension.
2. Phenotypic transition of aortic smooth muscle cells (SMC) has been demonstrated in atherosclerosis at single cell resolution.

### What new information does this article contribute?

1. Single cell multimodal profiling revealed persistent changes in the transcriptome and epigenome of vascular cells in aortas from diabetic mice despite glycemic control with the anti-diabetic sodium-glucose cotransporter 2 (SGLT2) inhibitor drug.
2. Diabetes induced reduction in the expression and chromatin accessibility of SMC contractile genes and increases in key fibrotic genes were not reversed by glucose normalization with the SGLT2 inhibitor.
3. Xenium spatial transcriptomic profiling of aortas from diabetic mice treated with and without the SGLT2 inhibitor also showed persistence of diabetes-induced changes in SMC contractile genes.

Diabetes is associated with vascular smooth muscle cell (SMC) dysfunction and accelerated vascular complications like atherosclerosis and hypertension. Such complications may persist even after glycemic control due to “metabolic memory” of prior hyperglycemia. To decipher the mechanisms, we used state-of-the-art single-cell (sc) multi-omics to examine the effect of glucose normalization (with the anti-diabetic SGLT2 inhibitor Dapagliflozin, DAPA) on transcriptomic and epigenomic changes associated with SMC phenotypic transition in type 2 diabetic db/db mice. We show contractile SMC population and SMC markers are decreased in diabetic mice and not reversed by DAPA. Conversely, fibromyocyte-like cells and key transcription factors regulating fibromyocyte phenotype are increased and remain increased after DAPA treatment. Furthermore, sc-ATAC-seq revealed key transcription factors driving contractile to fibromyocyte transition in diabetes and with DAPA. Spatial transcriptomics (Xenium) validated our scRNA-seq results. Human relevance was shown using scRNA-seq data from donor arteries, validating markers found in both db/db and db/dbDAPA mice. Together, our integrated datasets reveal diabetes/hyperglycemia-induced (epigenetic) memory in aortic SMC, as well as endothelial cells and fibroblasts with single cell/spatial resolution. Many changes are not reversed by an anti-diabetic drug, underscoring the unmet need for novel therapies to effectively target hyperglycemic memory and prevent long-term diabetic vascular complications.

